# Comparison of three quantitative approaches for estimating time-since-deposition from autofluorescence and morphological profiles of cell populations from forensic biological samples

**DOI:** 10.1101/2023.04.19.537512

**Authors:** Amanda Elswick Gentry, Sarah Ingram, M. Katherine Philpott, Kellie J. Archer, Christopher J. Ehrhardt

## Abstract

Determining when DNA recovered from a crime scene transferred from its biological source, i.e., a sample’s ‘time-since-deposition’ (TSD), can provide critical context for biological evidence. Yet, there remains no analytical techniques for TSD that are validated for forensic casework. In this study, we investigate whether morphological and autofluorescence measurements of forensically-relevant cell populations generated with Imaging Flow Cytometry (IFC) can be used to predict the TSD of ‘touch’ or trace biological samples. To this end, three different prediction frameworks for estimating the number of day(s) for TSD were evaluated: the elastic net, gradient boosting machines (GBM), and generalized linear mixed model (GLMM) LASSO. Additionally, we transformed these continuous predictions into a series of binary classifiers to evaluate the potential utility for forensic casework. Results showed that GBM and GLMM-LASSO showed the highest accuracy, with mean absolute error estimates in a hold-out test set of 29 and 21 days, respectively. Binary classifiers for these models correctly binned 94-96% and 98-99% of the age estimates as over/under 7 or 180 days, respectively. This suggests that predicted TSD using IFC measurements coupled to one or, possibly, a combination binary classification decision rules, may provide probative information for trace biological samples encountered during forensic casework.

## Introduction

An ongoing priority for forensic caseworking agencies has been developing signatures that can associate a time-since-deposition (TSD) with DNA evidence (OSAC pub, and review article). This can help determine whether a person of interest’s DNA was transferred within or outside of the time frame of the crime, which in turn informs its relevance. Many methods have been proposed for TSD, including Raman spectroscopy(1), infrared spectroscopy(2), mRNA signatures(3–5), bacterial profiling(6) and colorimetric assays(7). While promising, many of these techniques have been demonstrated on a limited range of TSDs (e.g., samples less than a month old). Further, the overwhelming majority of previously described methods have been applied to blood and/or saliva cells and have not been tested on ‘touch’ or trace biological samples which largely consist of keratinized epidermal cells and often comprise the majority of samples processed by DNA caseworking units.

One promising approach for determining TSD of trace biological samples focuses on Imaging Flow Cytometry (IFC) characterizations of epidermal cells recovered from the sample. Previous research has shown that morphological and/or autofluorescence profiles of cells derived from either buccal tissue or shed epidermal deposits change over time in mock evidentiary samples(8,9). Further, for shed epidermal cells, IFC profiles can be used to build predictive linear models that provide a probative time interval for sample deposition, e.g., whether the sample was deposited less than a week before collection, deposited between one week and two months, or deposited more than two months(9).

Using this as a foundation, the goal of this study was to test a different quantitative approach, based on continuous models in machine learning (ML), for predicting TSD from autofluorescence/morphological profiles of touch/trace cell populations. Continuous regression models have the potential to provide better resolved estimates of TSD compared to the categorical time intervals, e.g., TSD estimate of four days versus TSD estimate of less than a week. Previous work has applied ML to prediction problems in forensic science, including recent applications predicting tissue age and parameter selection for fluorescent molecular topography(10,11). ML has also been applied to IFC measurements to classify white blood cell types(12) as well as red blood cell types(13), to differentiate cancer cells from blood cells(14), and to predict gene expression from blood cells(15). To our knowledge, ML has never been applied to estimate TSD in epithelial cells from touch samples using IFC measurements.

While many forms of machine learning models could be applied to this prediction problem, of particular interest are those featuring an interpretable framework with the potential to yield biological insight into the mechanism by which measured features of collected cells change according to age. We chose to employ three distinct ML approaches: the elastic net using both the LASSO and the ridge regression penalties, gradient boosting machines (GBM), and generalized linear mixed model (GLMM) LASSO. The elastic net and GLMM models provide the greatest level of model interpretability while the GBM approach is more opaque but often provides superior prediction.

## Materials and Methods

### Sample collection

Trace epidermal cell samples were collected from participants following protocols approved by the Virginia Commonwealth University Institutional Review Board (VCU-IRB), protocol #HM20000454_CR9. Written informed consent was obt0061ined from all participants in this study. Epidermal samples were collected from 15 different individuals with some donors contributing multiple samples representing different time-since-depositions. This yielded a total of 47 different samples collected for the study. Sample deposition involved each donor holding a 50mL conical tube (Olympus Plastics, 28-108) using both hands for approximately three minutes. Tubes were then left at room temperature for designated periods of time ranging between 1-415 days.

Epidermal cells were collected from the substrate with two sterile cotton swabs (Puritan Inc., P/N 25-806 2WC), one prewetted with deionized water and the second swab dry. Cells were eluted off the swab by agitating 1.5mL of sterile 1xPBS for ∼10 minutes using a vortex. The resulting solution was filtered through 100µm filter paper into 1.5mL centrifuge tubes, and the samples were centrifuged at 21130xg for 10 minutes. The supernatant was pipetted from each tube until the remaining volume was between 50µL and 75µL. The pellet was resuspended by vortex agitation and stored at room temperature before flow cytometry.

### Imaging Flow Cytometry and data description

IFC was performed using an Amnis Imagestream X Mark II (Luminex Inc; Austin, TX, USA) equipped with 405nm, 488nm, 561nm, and 642nm lasers. Laser voltages were set at 120mW, 100mW, 100mW and 150mW, respectively. All events images were captured using five detector channels: 1 (430-505nm), 2 (505-560nm), 3 (560-595nm), 5 (640-745nm), and 6 (745-780nm). Channel 4 captured brightfield images. All images were captured at 40x magnification with enabled autofocus.

Morphological and autofluorescence measurements were extracted from cell populations by importing primary data files generated from IFC (i.e., ‘raw image files’; .rif) into IDEAS 6.2 Software (Luminex Inc; Austin, TX, USA). Next, intact cells were identified and differentiated from cell fragments, debris, or other non-biological material by selecting detected ‘events’ with areas greater than 1000µm and aspect ratios over 0.4 using Area_M04 x Aspect Ratio_M04 measurement variables respectively. The subpopulation of cell images was further filtered for in-focus images selecting cells with Gradient RMS values over 50 in the brightfield channel (i.e., non fluorescence channel).

Once each subpopulation were defined, data was extracted from individual cells using 14 different categories of morphological and autofluorescence measurements including area, aspect ratio, contrast, intensity, mean pixel, brightness detail intensity (‘R3’ pixel increment), brightness detail intensity (‘R7’ pixel increment), and compactness. Additionally, the ratio of Intensity and Brightness Detail Intensity measurements were calculated across several detector channels including Ch3:Ch1 (Brightness Detail Intensity R3), Ch3:Ch1 (Brightness Detail Intensity R7), Ch3:Ch1 (Intensity), Ch5:Ch1 (Intensity), Ch6:Ch1 (Intensity), and Ch6:Ch3 (Intensity). These yielded a total of 97 measurement variables for each detected cell. This primary data may be found under the title, “Imaging Flow Cytometry (IFC) dataset for cell populations recovered from ‘touch’/trace biological samples” at DOI: 10.6084/m9.figshare.22652344.

The observed cell dataset contained 97 measurements captured with IFC from each of 10,221 detected cell events, or “observations” (i.e., biological particles meeting the size, shape, and contrast criteria described above), across all the collected donor deposits. Samples were collected from 15 donors across multiple time points. In total, 47 different donor/timepoints (hereafter referred to as “samples”) are represented in the data. The number of timepoints samples varied by donor, with a maximum of 9 and a median count of 2. **Table 1** summarizes the number of observations collected per sample, at each timepoint. Minimum number of observations per donor/timepoint was 3, and maximum was 2,151, with a median of 78 observations. Observations were taken between 1 and 415 days since deposition, with a median of 35 days. We transformed raw time in days to time in ln(days) for the purpose of machine learning predictions.

**Table 1:**
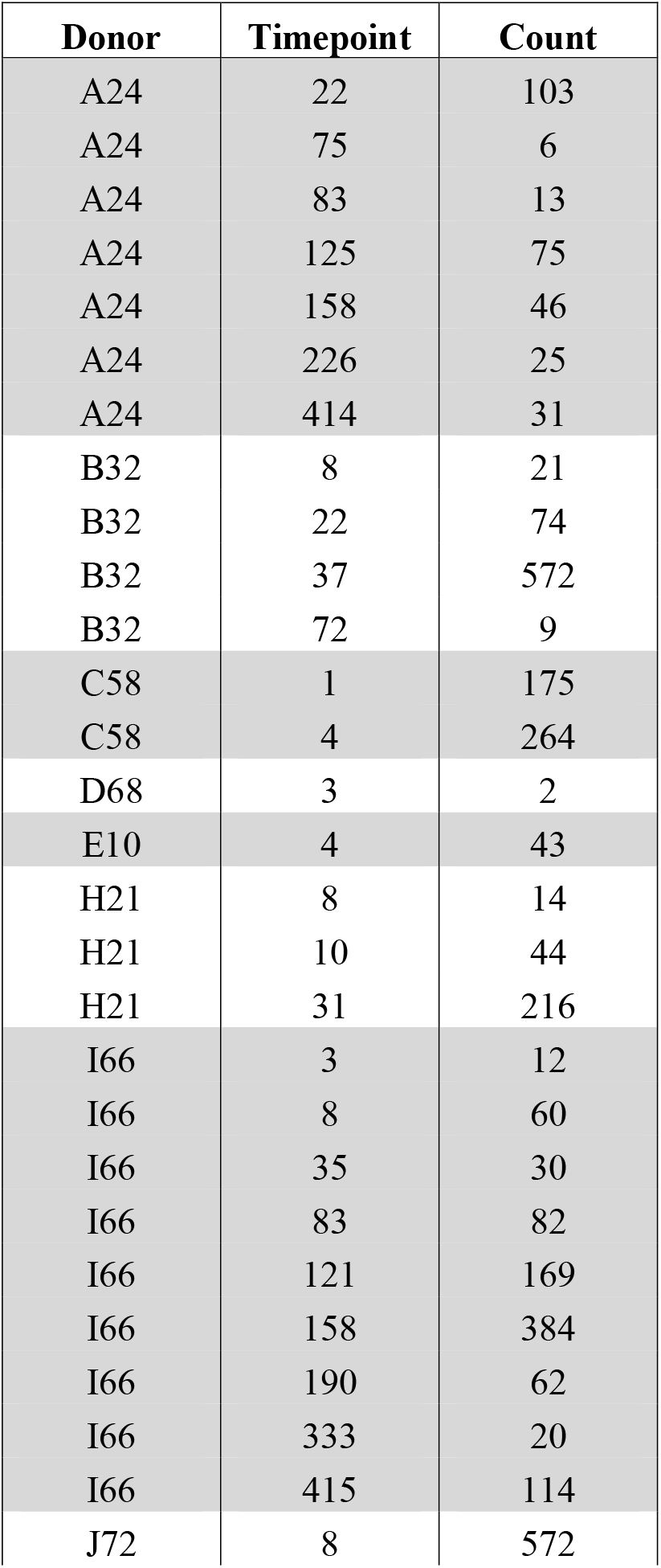
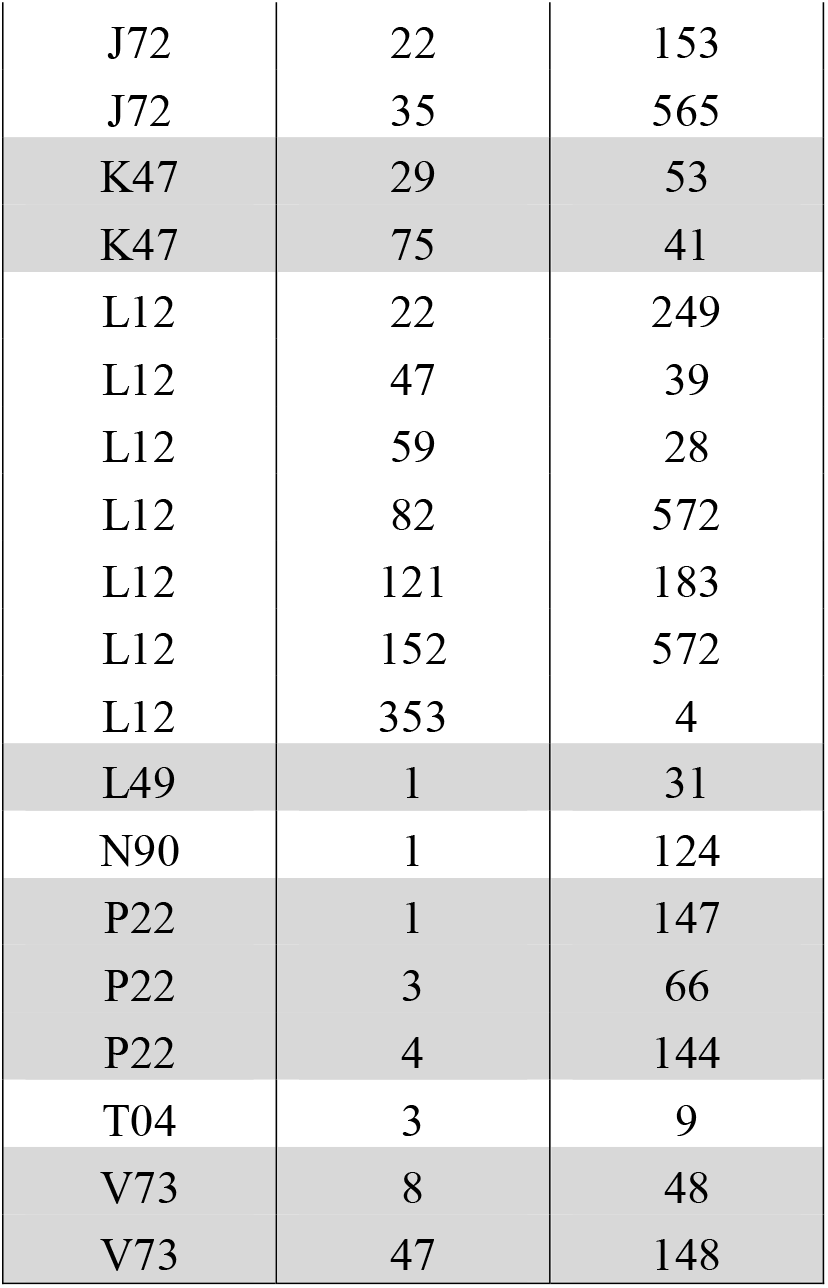
Number of observations taken from each donor at each timepoint.

### Training and test sets

Before applying any machine learning methods to predict time since deposition of touch samples, we divided our data into non-overlapping training and test sets. The test set represents a hold-out set kept in reserve for testing models constructed using the training set and contains points never used in training the models. The number of timepoints collected per donor, as well as the number of observations taken at each timepoint varied across donors. In order to balance the test and training sets, we reserved 20% of the observations from each sample for the test set. We rounded the number of observations in the test set up where 20% of the number of observations did not result in a whole number. Furthermore, in cases where the number of observations in the training set for a given sample exceeded the 90th percentile of observation counts across all donors/timepoints, we capped the number of observations in the training set at the 90th percentile and reserved the remainder for the test set. **Supplemental Table S1** summarizes the number of observations in the test and training sets for each sample.

### Machine learning models

We applied 3 separate supervised machine learning paradigms to predict the time since sample deposition in ln(days). First, we applied the elastic net under two conditions, with the alpha mixing parameter at 1 for the LASSO fit and with alpha at 0 for ridge regression(16). Second, we applied Gradient Boosting Machines (GBM)(17), and third, we applied the generalized linear mixed model LASSO(18). Data were analyzed in R (version 4.1.1) using the glmnet (version 4.1.4)(19), gbm (version 2.1.8)(20), and glmmLasso (version 1.6.2)(18) packages. All scripts used for data processing and analysis have been deposited into a public GitHub repository available here. [https://github.com/AEGentry/ForensicSci]

Each of our applied ML approaches requires cross-validation to select the optimal tuning and/or hyperparameters. After cross-validation identifies the optimal model for each approach, a final model it fit to the training data and assessed. For the LASSO and ridge regression, we fit the models to the training data using 10-fold cross validation to choose the shrinkage parameter according to the built-in cross-validation functionality of the cv.glmnet command. The GBM fitting utilized 10-fold cross-validation to tune the model and assess a grid of potential hyperparameters. We tested potential hyperparameters for shrinkage (0.001, 0.01, and 0.05), interaction depth (1 and 2), and minimum observations per node (5, 10, and 15). We chose the optimal GBM model as the one which produced the minimum average mean squared error (MSE) across the folds. Once the optimal tuning parameters have been chosen, the final model fits the training data using the recommended 0.5 bag fraction and chooses the optimal number of trees (ie., iterations) via 10-fold cross-validation., For the GLMM models, the glmmLasso package does not include built-in fitting via cross-validation, so we fit the model to the training data using 10-fold cross-validation to choose the shrinkage parameter across values ranging from 5 to 100, in increments of 5.

### Assessing performance

To assess prediction performance and compare predictions across models, we calculated mean squared error (MSE) and mean absolute error (MAE) between the observed and predicted days since deposition, both on the ln scale and the original scale, within the test set. Additionally, for a secondary, statistical sensitivity analysis, we used N-fold cross-validation whereby we applied each ML model to a subset of the training data 47 times, each time holding out one of the 47 sample cell populations and then used the hold-out donor/timepoint cell population for prediction. These donor/timepoint sample populations are described in **Supplemental Table S1**. We also calculated MSE and MAE for the predictions within each of these donor/timepoint hold-out sets and summarized the means of these measures across the 47 hold out sets.

Each of the ML approaches predicts time since deposition as a continuous value (either ln(days) or days). To demonstrate the potential real-world usefulness of this prediction paradigm, we categorized these continuous predictions, both in the test set and within the N-fold cross-validation sample hold-out sets. We categorized the predictions using six binary schemes for time since deposition: less than or greater than 7 days, 30 days, 60 days, 90 days, 120 days, or 180 days, and summarized the performance of each using the proportion of predictions properly categorized by each binary cutoff.

## Results

### Model fitting

The full dataset containing 10,221 observations was divided into a training set with 6,414 observations, or approximately 62.8% of the data, and a test set with 3,807 observations, or approximately 37.8% of the data. The lambda penalty term minimizing cross-validation error was 0.00081 and 0.05997 in the LASSO and ridge models, respectively; the LASSO model retained 83 predictors with non-zero coefficients. The GBM hyperparameters achieving best fit utilized shrinkage=0.05, a minimum number of observations per node of 15, interaction depth of 2, and an optimal number of trees of 5786. For the GLMM model, the lambda penalty term minimizing cross-validation error was 90 and that model retained 94 predictors with non-zero coefficients.

### Prediction performance for estimating TSD in days

Overall MAE for ln(days) and the original scale days for all four models are detailed in **Table 2** for the test sets. MSE and training set error are reported in **Supplemental Table S2**. Errors on the original scale were estimated by transforming the predicted ln(days) to the original scale and calculating MSE and MAE on the transformed values. Optimal models are fit using MSE of ln(days) internal to the ML algorithms, as this loss function optimizes prediction, while MAE in raw days is useful for reporting performance as it is the more readily interpretable, intuitive metric. Within the training data, the GBM model outperformed all other models, both in MSE and MAE, although performance in the GLMM model was similar (e.g., 17.2 MAE days for GBM and 21.8 MAE days for GLMM). In the hold-out test set, the GLMM model out performed the others, with a MSE of 0.18 and a MAE of 0.31, while the GBM model performed nearly as well, with a MSE of 0.33 and a MAE of 0.41.

**Table 2:**
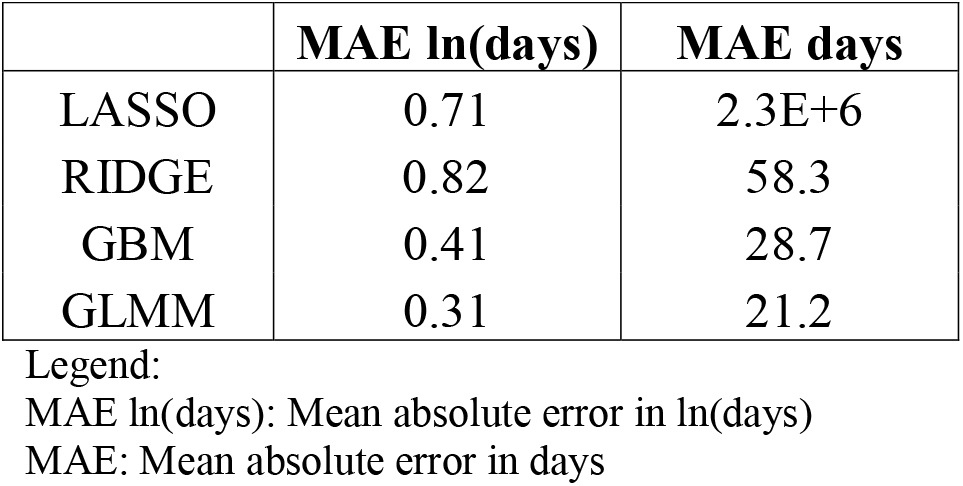
Mean absolute prediction error in the test set.

In the sensitivity analysis that used N-fold cross-validation for each donor cell population (**Table 3** and **Supplemental Table S3**), the GBM model again outperformed all other models with average MSE of 1.28 and MAE of 0.82, while the GLMM model achieved average MSE of 1.41 and MAE of 0.89. **Figs 1 and 2** illustrate the absolute value of the prediction error in days for the GBM and GLMM models in the hold-out test set, respectively. The median absolute error was 17.3 (GBM) and 12.8 (GLMM) days with third quartiles only reaching 37.8 (GBM) and 25.7 (GLMM) days. Notably, while the absolute error does increase as the true TSD increases, the majority of samples, particularly those that were less than two months old, error was extremely low.

**Table 3:** Mean absolute prediction error in the N-fold cross-validation set.

|  | **MAE ln(days)** | **MAE days** |
| --- | --- | --- |
| LASSO | 0.99 | 743.4 |
| RIDGE | 1.05 | 281.7 |
| GBM | 0.82 | 46 |
| GLMM | 0.89 | 46.8 |

MAE ln(days): Mean absolute error in ln(days)
MAE days: Mean absolute error in days

**Figs 1-2:**
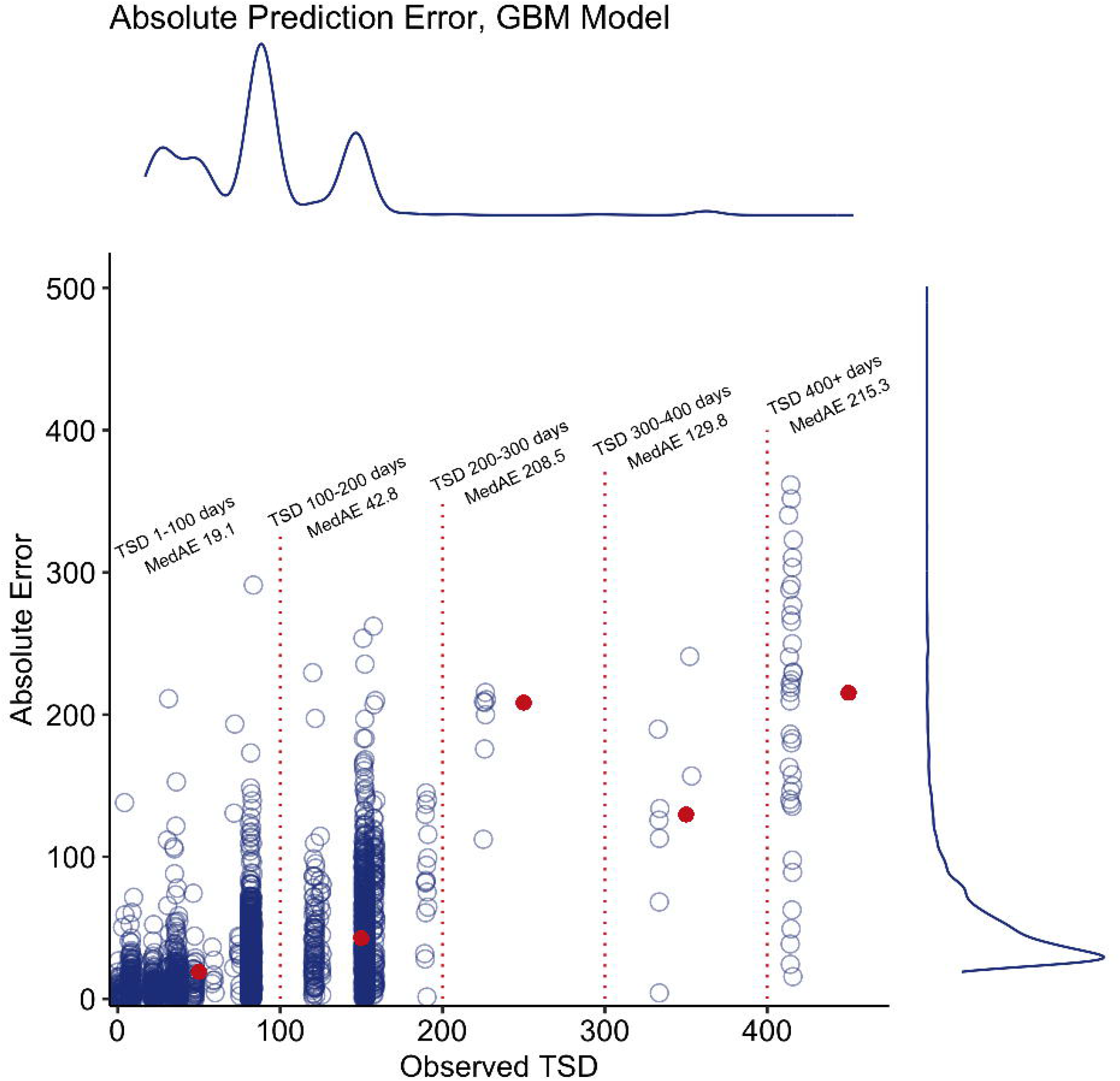
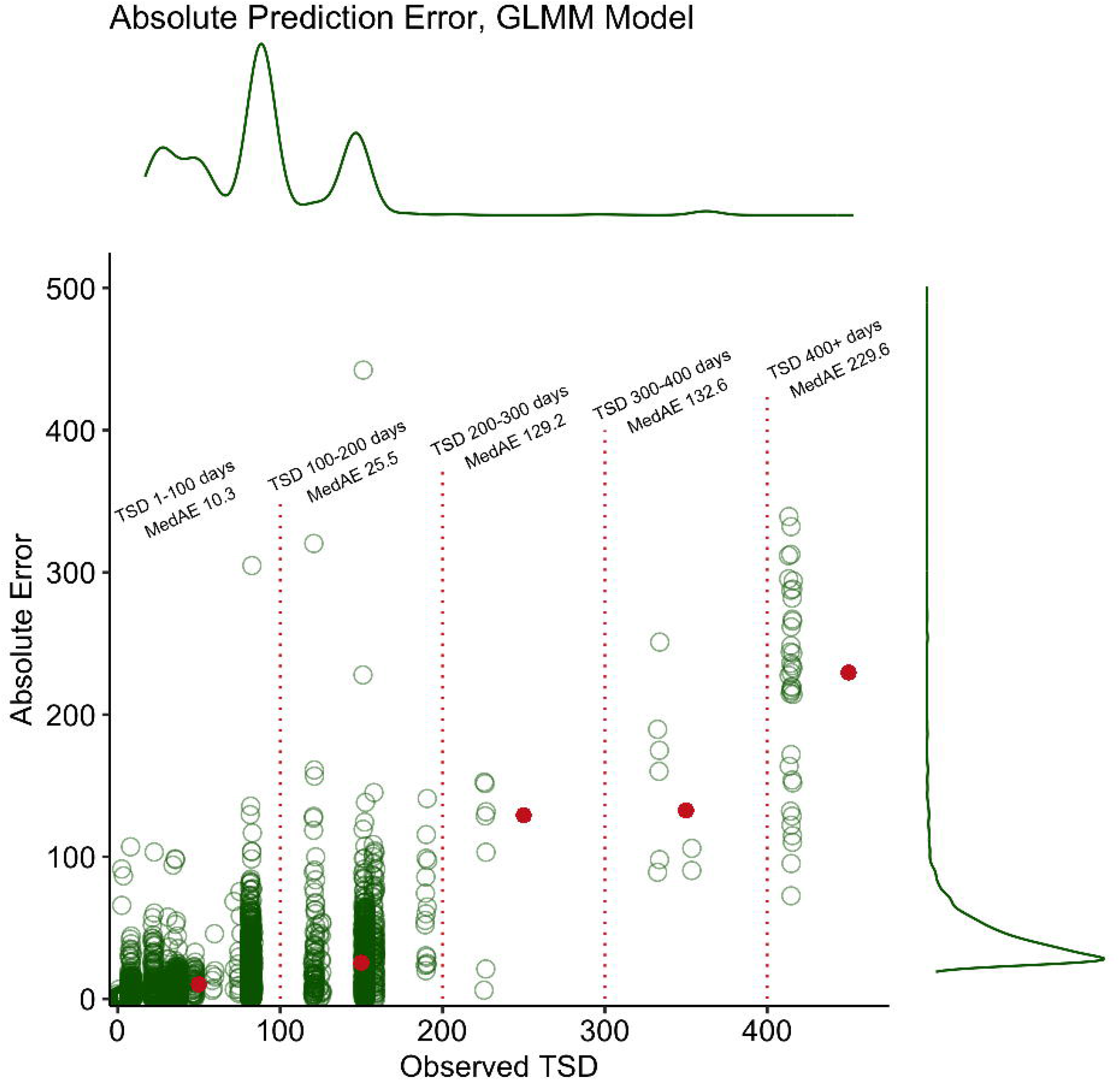
Absolute prediction error scatterplots with density curves for GBM (Fig 1) and GLMM (Fig 2) models. Median absolute error for various bins shown in red points.

This trend is not unexpected given the structural and biochemical heterogeneity of shed epidermal cells within touch/trace samples(21). Indeed, given the established variability within an epidermal cell population deposited by one person at a single timepoint, it is not unreasonable to expect that morphological and/or autofluorescence changes occurring as the sample ages to increasingly diverge over time, and for subsets of the cells to change at different rates.

### Prediction performance for estimating TSD with binary time intervals

TSD estimates from each regression model were also used to classify each donor cell population into one of two time intervals that span the entire observation period for this study (i.e., between 0 and 415 days). To further explore the potential utility for forensic casework of TSD estimates, we chose a series of binary cutoff points to use as classifiers. Using this approach, we simply transformed the model predicted TSD in days and calculated the proportion of these estimates which correctly classified the TSD as older or newer than a given cutoff. We did not conduct any new classifying procedures, rather this reporting simply represents a categorization of the continuous prediction.

The proportions of observations properly classified at different timepoints according to the series of binary cutoffs are shown in aggregate for the test set and the donor/timepoint set in **Table 4**. In the test set, the GBM and GLMM models performed similarly, with the models correctly classifying observations between 81-99% of the time. Performance was similar in the donor/timepoint set, with the GBM and GLMM models properly classifying observations between 70-97% of the time. In both sets, the best prediction occurred at the tails, that is, for the lowest binary classifier (greater/less than one week) and the highest (greater/less than 180 days.) For the GBM and GLMM models, the best performing approaches, we also report the breakdown of classifier performance by donor (**Supplemental Tables S4-S5**, respectively) and by donor/timepoint (**Supplemental Tables S6-S7**, respectively.) Overall, these summaries do not indicate any extreme donor or donor/timepoint outliers, although samples with more observations tended to perform better.

**Table 4:** Proportions of properly classified timepoints using a series of binary cutoff values for the hold-out test set and the donor/timepoint set.

|  | **LASSO** |  | **Ridge** |  | **GBM** |  | **GLMM** |  | **Total** |  |
| --- | --- | --- | --- | --- | --- | --- | --- | --- | --- | --- |
|  | *Test* | *DT* | *Test* | *DT* | *Test* | *DT* | *Test* | *DT* | *Test* | *DT* |
| **Under 7** | 0.973 | 0.932 | 0.961 | 0.897 | 0.983 | 0.964 | 0.996 | 0.969 | 259 | 1017 |
| **Under 30** | 0.787 | 0.694 | 0.724 | 0.668 | 0.915 | 0.756 | 0.913 | 0.701 | 755 | 2408 |
| **Under 60** | 0.615 | 0.79 | 0.548 | 0.771 | 0.834 | 0.825 | 0.919 | 0.919 | 1222 | 4006 |
| **Under 90** | 0.872 | 0.849 | 0.847 | 0.823 | 0.811 | 0.845 | 0.847 | 0.915 | 2842 | 4729 |
| **Under 120** | 0.843 | 0.820 | 0.807 | 0.795 | 0.842 | 0.816 | 0.891 | 0.869 | 2842 | 4729 |
| **Under 180** | 0.940 | 0.936 | 0.965 | 0.95 | 0.935 | 0.947 | 0.961 | 0.945 | 3739 | 6158 |

Test: hold-out test set
DT: donor/timepoint hold out sets

## Discussion

The results of this study demonstrate the potential for machine learning-based methods to provide probative TSD estimates of ‘touch’ biological samples from IFC measurements of cell populations. We tested four separate models across three different machine learning paradigms for prediction and found that GBM and GLMM models outperformed both the LASSO and the ridge regression implementations of the elastic net. This disparity in performance is not wholly unexpected, as ensemble learners such as GBMs often outperform penalized regressions (ie elastic net) in some data, while we postulate that improved performance in the GLMM models was on account of the random effects in that model which take inter-donor covariance into account. The best performing model (GLMM) showed an overall MAE rate of ∼21 days for predicted time since deposition of blinded (or holdout) cell population samples.

While this level of uncertainty may seem large for forensic applications, we noted that time discrepancies varied with the TSD of the sample. In particulare, cell populations deposited less than one week previously had MAE of 1.1 days (GLMM model), whereas cell populations deposited more than 6 months previously had MAE of 161.9 days. Within a forensic context, because it is relatively common for the prosecution and defense to theorize highly disparate time frames for DNA deposition, these TSD estimates can be quite probative. One side may propose that DNA recovered from a crime scene was deposited at the time of the crime, and the other may claim it was transferred days, weeks, months or years before or after the crime event(22–24). As such, even broad TSD estimates may be useful for both forensic laboratories and the legal system in distinguishing relevant from misleading biological material in the course of a criminal investigation.

To investigate how TSD estimates could be applied within this context, we considered an alternative framework for interpreting TSD estimates using a series of binary time intervals; determining whether a sample was deposited greater/less than 7, 30, 60, 90, 120, or 180 days. With this scheme, both GBM and GLMM models were able to properly classify the age of a sample most accurately at the extremes of the TSD range, with over 99% accuracy for categorizing samples as greater/less than one week old and 96% accuracy for categorizing samples as greater/less than 180 days old in the GLMM models predicting in the test set. We further demonstrated that this performance remained strong in the sensitivity analysis testing the model’s ability to predict TSD for a blind sample, i.e. one that was held out, to which the training model was naive. In this set of held-out samples, the GLMM model’s overall classification accuracy was 97% for categorizing a sample as 7 days old or less and 95% accurate for categorizing as 180 days old or less, in the GLMM model. As data clusters towards the center of the distribution, the ability of the model to delineate properly between classes diminishes slightly with observed accuracies of 91%, 92%, 85%, and 90% for categorizing age of cells as greater/less than 30, 60, 90, and 120 days, respectively.

Several important limitations of note include the moderate sample size employed here. We had 15 independent donors who provided samples that were subsequently measured at multiple timepoints. A related limitation relates to the uneven number of observations taken from each subject, as well as the highly varying nature of the timepoints. We expect that results from this proof-of-concept study could serve as the basis for a formal validation effort that tests the robustness of TSD estimation on a larger donor set and, possibly, conditions that can be encountered by caseworking agencies (e.g., mixture samples).

Lastly, we selected only four, largely regression-centric machine learning models to test here. This limitation, however, lends itself to important implications for practice. These four machine learning models were selected explicitly because they were non-”black box” in nature, allowing the user to understand the manner in which each independent variable contributes to the ultimate prediction. Such models can be more intuitive and interpretable compared to other computational approaches, such as deep learning which may facilitate its adoption by the forensic science community. Furthermore, we note that our approaches employed continuous regression models. Evaluation of the error structure from these models may indicate that a discrete response model may be warranted. Future work aims to evaluate the utility of such models for this and similar data.

As a final note, the field should consider a change in nomenclature from the “time since deposition” language found in the existing literature, which may be out of step with our current understanding of the realities of direct and indirect transfer of biological material. Because these methods are based on cellular changes over time, these methods are actually predicting time since primary transfer from the biological source. Sometimes this is the same as time since deposition (e.g. if a person touches an object and their cells transfer), but if indirect transfer takes place, it may not. In other words, if cells are transferred from their biological source to one substrate, and then from that substrate to an item at the crime scene at a different point in time— a phenomenon that has been shown to occur in both studies and real life(23,25) - the time predicted should theoretically be associated with the original, or “primary” transfer event (and this would be desired).

## Supporting Information

**S1 Table: Number of observations in the hold-out and training sets for each donor/timepoint**.

Legend:

Samples By Time: Donor/timepoint combination

N Observations: Total number of observations for a given donor/timepoint

N Test: Number of observations in the hold-out test set

N Train: Number of observations in the training set

**S2 Table: Mean squared and mean absolute prediction errors for test and training sets**.

Legend:

MSE ln(days) Train: Mean squared error in ln(days) in the training set

MSE days Train: Mean squared error in days in the training set

MSE ln(days) Test: Mean squared error in ln(days) in the test set

MSE days Test: Mean squared error in days in the test set

MAE ln(days) Train: Mean absolute error in ln(days) in the training set

MAE days: Mean absolute error in days in the training set

**Supplemental Table S3: Mean squared prediction error in the N-fold cross-validation set**.

Legend:

MSE ln(days): Mean squared error in ln(days)

MSE days: Mean squared error in days

**Supplemental Table S4: Classifier performance for each hold-out donor cell population, using GBM models**.

Legend:

Count: Number of cells that properly classify

Prop: Proportion of cells that properly classify

Total: Total cell count for the donor population

**Supplemental Table S5: Classifier performance for each hold-out donor cell population, using GLMM models**.

Legend:

Count: Number of cells that properly classify

Prop: Proportion of cells that properly classify

Total: Total cell count for the donor population

**Supplemental Table S6: Classifier performance for each hold-out donor/timepoint cell population, using GBM models**.

Legend:

Count: Number of cells that properly classify

Prop: Proportion of cells that properly classify

Total: Total cell count for the donor population

**Supplemental Table S7: Classifier performance for each hold-out donor/timepoint cell population, using GLMM models**.

Legend:

Count: Number of cells that properly classify

Prop: Proportion of cells that properly classify

Total: Total cell count for the donor population

## Notes

### Competing Interest Statement

The authors have declared no competing interest.

## References

1. Doty KC, McLaughlin G, Lednev IK. A Raman “spectroscopic clock” for bloodstain age determination: the first week after deposition. Anal Bioanal Chem. 2016 Jun;408(15):3993–4001.

2. Edelman G, Manti V, van Ruth SM, van Leeuwen T, Aalders M. Identification and age estimation of blood stains on colored backgrounds by near infrared spectroscopy. Forensic Sci Int. 2012 Jul 10;220(1–3):239–44.

3. Bauer M, Polzin S, Patzelt D. Quantification of RNA degradation by semi-quantitative duplex and competitive RT-PCR: a possible indicator of the age of bloodstains? Forensic Sci Int. 2003 Dec 17;138(1–3):94–103.

4. Anderson S, Howard B, Hobbs GR, Bishop CP. A method for determining the age of a bloodstain. Forensic Sci Int. 2005 Feb 10;148(1):37–45.

5. Salzmann AP, Russo G, Kreutzer S, Haas C. Degradation of human mRNA transcripts over time as an indicator of the time since deposition (TsD) in biological crime scene traces. Forensic Sci Int Genet. 2021 Jul;53:102524.

6. Díez López C, Kayser M, Vidaki A. Estimating the Time Since Deposition of Saliva Stains With a Targeted Bacterial DNA Approach: A Proof-of-Principle Study. Front Microbiol. 2021 Jun 2;12:647933.

7. Shin J, Choi S, Yang J-S, Song J, Choi J-S, Jung H-I. Smart Forensic Phone: Colorimetric analysis of a bloodstain for age estimation using a smartphone. Sens Actuators B Chem. 2017 May 1;243:221–5.

8. Brocato ER, Philpott MK, Connon CC, Ehrhardt CJ. Rapid differentiation of epithelial cell types in aged biological samples using autofluorescence and morphological signatures. PLoS One. 2018 May 17;13(5):e0197701.

9. Ingram S, Philpott MK, Ehrhardt CJ. Novel cellular signatures for determining time since deposition for trace DNA evidence. Forensic Science International: Genetics Supplement Series. 2022 Dec 1;8:268–70.

10. Alsaleh H, McCallum NA, Halligan DL, Haddrill PR. A multi-tissue age prediction model based on DNA methylation analysis. Forensic Science International: Genetics Supplement Series. 2017 Dec 1;6:e62–4.

11. Wang H, Bian C, Kong L, An Y, Du Y, Tian J. A Novel Adaptive Parameter Search Elastic Net Method for Fluorescent Molecular Tomography. IEEE Trans Med Imaging. 2021 May;40(5):1484–98.

12. Lippeveld M, Knill C, Ladlow E, Fuller A, Michaelis LJ, Saeys Y, et al. Classification of Human White Blood Cells Using Machine Learning for Stain-Free Imaging Flow Cytometry. Cytometry A. 2020 Mar;97(3):308–19.

13. Demagny J, Roussel C, Le Guyader M, Guiheneuf E, Harrivel V, Boyer T, et al. Combining imaging flow cytometry and machine learning for high-throughput schistocyte quantification: A SVM classifier development and external validation cohort. EBioMedicine. 2022 Sep;83:104209.

14. Nissim N, Dudaie M, Barnea I, Shaked NT. Real-Time Stain-Free Classification of Cancer Cells and Blood Cells Using Interferometric Phase Microscopy and Machine Learning. Cytometry A. 2021 May;99(5):511–23.

15. Chlis N-K, Rausch L, Brocker T, Kranich J, Theis FJ. Predicting single-cell gene expression profiles of imaging flow cytometry data with machine learning. Nucleic Acids Res. 2020 Nov 18;48(20):11335–46.

16. Zou H, Hastie T. Regularization and variable selection via the elastic net. J R Stat Soc Series B Stat Methodol. 04/2005;67(2):301–20.

17. Friedman JH. Greedy function approximation: A gradient boosting machine. Ann Stat. 10/2001;29(5):1189–232.

18. Groll A, Tutz G. Variable selection for generalized linear mixed models by L 1-penalized estimation. Stat Comput. 2014 Mar;24(2):137–54.

19. Friedman Jerome, Hastie T, Simon N, Qian J, Tibshirani R. glmnet: Lasso and Elastic-Net Regularized Generalized Linear Models [Internet]. Available from: https://cran.r-project.org/web/packages/glmnet/index.html

20. Bühlmann P, Hothorn T. Boosting Algorithms: Regularization, Prediction and Model Fitting. SSO Schweiz Monatsschr Zahnheilkd. 2007 Nov;22(4):477–505.

21. Burrill J, Daniel B, Frascione N. A review of trace “Touch DNA” deposits: Variability factors and an exploration of cellular composition. Forensic Sci Int Genet. 2019 Mar;39:8–18.

22. Elster N. How Forensic DNA Evidence Can Lead to Wrongful Convictions. JSTOR Daily [Internet]. 2017 Dec 6; Available from: https://daily.jstor.org/forensic-dna-evidence-can-lead-wrongful-convictions/

23. Murphy EE. How DNA Evidence Incriminated an Impossible Suspect. 2015 Oct 26; Available from: https://newrepublic.com/article/123177/how-dna-evidence-incriminated-impossible-suspect

24. Jabali-Nash N. Chandra Levy Murder Trial: DNA on Victim Doesn’t Match Defendant, Expert Testifies. 2010 Nov 4; Available from: https://www.cbsnews.com/news/chandra-levy-murder-trial-dna-on-victim-doesnt-match-defendant-expert-testifies/

25. van Oorschot RAH, Szkuta B, Meakin GE, Kokshoorn B, Goray M. DNA transfer in forensic science: A review. Forensic Sci Int Genet. 2019 Jan;38:140–66.

