## Supplemental Table 1 for "Comparison of three quantitative approaches for estimating time-since-deposition from autofluorescence and morphological profiles of cell populations from forensic biological samples"

**Table S1: Number of observations in the hold-out and training sets for each donor/timepoint.**

| **Sample By Time** | **N Observations** | **N Test** | **N Train** |
| --- | --- | --- | --- |
| A24.125 | 94 | 19 | 75 |
| A24.158 | 58 | 12 | 46 |
| A24.22 | 129 | 26 | 103 |
| A24.226 | 32 | 7 | 25 |
| A24.414 | 39 | 8 | 31 |
| A24.75 | 8 | 2 | 6 |
| A24.83 | 17 | 4 | 13 |
| B32.22 | 93 | 19 | 74 |
| B32.37 | 779 | 207 | 572 |
| B32.72 | 12 | 3 | 9 |
| B32.8 | 27 | 6 | 21 |
| C58.1 | 219 | 44 | 175 |
| C58.4 | 331 | 67 | 264 |
| D68.3 | 3 | 1 | 2 |
| E10.4 | 54 | 11 | 43 |
| H21.10 | 56 | 12 | 44 |
| H21.31 | 271 | 55 | 216 |
| H21.8 | 18 | 4 | 14 |
| I66.121 | 212 | 43 | 169 |
| I66.158 | 481 | 97 | 384 |
| I66.190 | 78 | 16 | 62 |
| I66.3 | 15 | 3 | 12 |
| I66.333 | 26 | 6 | 20 |
| I66.35 | 38 | 8 | 30 |
| I66.415 | 143 | 29 | 114 |
| I66.8 | 76 | 16 | 60 |
| I66.83 | 103 | 21 | 82 |
| J72.22 | 192 | 39 | 153 |
| J72.35 | 707 | 142 | 565 |
| J72.8 | 856 | 284 | 572 |
| K47.29 | 67 | 14 | 53 |
| K47.75 | 52 | 11 | 41 |
| L12.121 | 229 | 46 | 183 |
| L12.152 | 1252 | 680 | 572 |
| L12.22 | 312 | 63 | 249 |
| L12.353 | 6 | 2 | 4 |
| L12.47 | 49 | 10 | 39 |
| L12.59 | 35 | 7 | 28 |
| L12.82 | 2151 | 1579 | 572 |
| L49.1 | 39 | 8 | 31 |
| N90.1 | 156 | 32 | 124 |
| P22.1 | 184 | 37 | 147 |
| P22.3 | 83 | 17 | 66 |
| P22.4 | 180 | 36 | 144 |
| T04.3 | 12 | 3 | 9 |
| V73.47 | 186 | 38 | 148 |
| V73.8 | 61 | 13 | 48 |

Legend:

Samples By Time: Donor/timepoint combination

N Observations: Total number of observations for a given donor/timepoint

N Test: Number of observations in the hold-out test set

N Train: Number of observations in the training set
