## Supplemental Table 2 for "Comparison of three quantitative approaches for estimating time-since-deposition from autofluorescence and morphological profiles of cell populations from forensic biological samples"

**Table S2: Mean squared and mean absolute prediction errors for test and training sets.**

|  | **MSE ln(days)**  **Train** | **MSE**  **days**  **Train** | **MSE ln(days)**  **Test** | **MSE**  **days**  **Test** | **MAE ln(days)**  **Train** | **MAE**  **days**  **Train** |
| --- | --- | --- | --- | --- | --- | --- |
| LASSO | 0.79 | 7334.4 | 0.93 | 2.081E+16 | 0.68 | 37.6 |
| RIDGE | 0.95 | 21406.6 | 0.99 | 226057.9 | 0.76 | 40.4 |
| GBM | 0.14 | 1134.2 | 0.33 | 2256.4 | 0.29 | 17.2 |
| GLMM | 0.27 | 2040.9 | 0.18 | 2329.5 | 0.39 | 21.8 |

Legend:

MSE ln(days) Train: Mean squared error in ln(days) in the training set

MSE days Train: Mean squared error in days in the training set

MSE ln(days) Test: Mean squared error in ln(days) in the test set

MSE days Test: Mean squared error in days in the test set

MAE ln(days) Train: Mean absolute error in ln(days) in the training set

MAE days: Mean absolute error in days in the training set
