## Supplemental Table 3 for "Comparison of three quantitative approaches for estimating time-since-deposition from autofluorescence and morphological profiles of cell populations from forensic biological samples"

**Supplemental Table S3: Mean squared prediction error in the N-fold cross-validation set.**

|  | **MSE ln(days)** | **MSE days** |
| --- | --- | --- |
| LASSO | 1.67 | 1.2E+10 |
| RIDGE | 1.77 | 2.7E+8 |
| GBM | 1.28 | 6995.2 |
| GLMM | 1.41 | 9561.9 |

Legend:

MSE ln(days): Mean squared error in ln(days)

MSE days: Mean squared error in days
