## Supplemental Table 4 for "Comparison of three quantitative approaches for estimating time-since-deposition from autofluorescence and morphological profiles of cell populations from forensic biological samples"

**Supplemental Table S4: Classifier performance for each hold-out donor cell population, using GBM models.**

|  | **Under 7 days** | | **Under 30 days** | | **Under 60 days** | | **Under 90 days** | | **Under 120 days** | | **Under 180 days** | |  |
| --- | --- | --- | --- | --- | --- | --- | --- | --- | --- | --- | --- | --- | --- |
| **Donor** | *Count* | *Prop* | *Count* | *Prop* | *Count* | *Prop* | *Count* | *Prop* | *Count* | *Prop* | *Count* | *Prop* | **Total** |
| A24 | 295 | 0.99 | 202 | 0.68 | 163 | 0.55 | 145 | 0.48 | 140 | 0.47 | 248 | 0.83 | 299 |
| B32 | 669 | 0.99 | 361 | 0.53 | 607 | 0.9 | 650 | 0.96 | 663 | 0.98 | 671 | 0.99 | 676 |
| C58 | 332 | 0.76 | 438 | 1 | 439 | 1 | 439 | 1 | 439 | 1 | 439 | 1 | 439 |
| D68 | 1 | 0.5 | 1 | 0.5 | 2 | 1 | 2 | 1 | 2 | 1 | 2 | 1 | 2 |
| E10 | 18 | 0.42 | 41 | 0.95 | 43 | 1 | 43 | 1 | 43 | 1 | 43 | 1 | 43 |
| H21 | 251 | 0.92 | 85 | 0.31 | 258 | 0.94 | 269 | 0.98 | 272 | 0.99 | 272 | 0.99 | 274 |
| I66 | 925 | 0.99 | 850 | 0.91 | 717 | 0.77 | 588 | 0.63 | 510 | 0.55 | 665 | 0.71 | 933 |
| J72 | 1272 | 0.99 | 622 | 0.48 | 1189 | 0.92 | 1274 | 0.99 | 1284 | 1 | 1290 | 1 | 1290 |
| K47 | 94 | 1 | 79 | 0.84 | 76 | 0.81 | 80 | 0.85 | 91 | 0.97 | 94 | 1 | 94 |
| L12 | 1617 | 0.98 | 1140 | 0.69 | 891 | 0.54 | 1253 | 0.76 | 1102 | 0.67 | 1566 | 0.95 | 1647 |
| L49 | 12 | 0.39 | 28 | 0.9 | 30 | 0.97 | 30 | 0.97 | 30 | 0.97 | 30 | 0.97 | 31 |
| N90 | 23 | 0.19 | 108 | 0.87 | 121 | 0.98 | 121 | 0.98 | 122 | 0.98 | 122 | 0.98 | 124 |
| P22 | 271 | 0.76 | 354 | 0.99 | 356 | 1 | 356 | 1 | 356 | 1 | 356 | 1 | 357 |
| T04 | 6 | 0.67 | 8 | 0.89 | 9 | 1 | 9 | 1 | 9 | 1 | 9 | 1 | 9 |
| V73 | 192 | 0.98 | 134 | 0.68 | 163 | 0.83 | 188 | 0.96 | 194 | 0.99 | 195 | 0.99 | 196 |

Legend:

Count: Number of cells that properly classify

Prop: Proportion of cells that properly classify

Total: Total cell count for the donor population
