## Supplemental Table 5 for "Comparison of three quantitative approaches for estimating time-since-deposition from autofluorescence and morphological profiles of cell populations from forensic biological samples"

**Supplemental Table S5: Classifier performance for each hold-out donor cell population, using GLMM models.**

|  | **Under 7 days** | | **Under 30 days** | | **Under 60 days** | | **Under 90 days** | | **Under 120 days** | | **Under 180 days** | |  |
| --- | --- | --- | --- | --- | --- | --- | --- | --- | --- | --- | --- | --- | --- |
| **Donor** | *Count* | *Prop* | *Count* | *Prop* | *Count* | *Prop* | *Count* | *Prop* | *Count* | *Prop* | *Count* | *Prop* | **Total** |
| A24 | 295 | 0.99 | 191 | 0.64 | 142 | 0.47 | 139 | 0.46 | 134 | 0.45 | 246 | 0.82 | 299 |
| B32 | 671 | 0.99 | 361 | 0.53 | 625 | 0.92 | 662 | 0.98 | 671 | 0.99 | 675 | 1 | 676 |
| C58 | 196 | 0.45 | 433 | 0.99 | 438 | 1 | 438 | 1 | 438 | 1 | 439 | 1 | 439 |
| D68 | 1 | 0.5 | 1 | 0.5 | 2 | 1 | 2 | 1 | 2 | 1 | 2 | 1 | 2 |
| E10 | 10 | 0.23 | 36 | 0.84 | 42 | 0.98 | 43 | 1 | 43 | 1 | 43 | 1 | 43 |
| H21 | 265 | 0.97 | 90 | 0.33 | 263 | 0.96 | 270 | 0.99 | 272 | 0.99 | 273 | 1 | 274 |
| I66 | 926 | 0.99 | 844 | 0.9 | 680 | 0.73 | 556 | 0.6 | 456 | 0.49 | 713 | 0.76 | 933 |
| J72 | 1279 | 0.99 | 593 | 0.46 | 1233 | 0.96 | 1280 | 0.99 | 1285 | 1 | 1289 | 1 | 1290 |
| K47 | 94 | 1 | 74 | 0.79 | 70 | 0.74 | 87 | 0.93 | 92 | 0.98 | 93 | 0.99 | 94 |
| L12 | 1625 | 0.99 | 1052 | 0.64 | 757 | 0.46 | 1093 | 0.66 | 995 | 0.6 | 1604 | 0.97 | 1647 |
| L49 | 9 | 0.29 | 27 | 0.87 | 31 | 1 | 31 | 1 | 31 | 1 | 31 | 1 | 31 |
| N90 | 16 | 0.13 | 109 | 0.88 | 116 | 0.94 | 121 | 0.98 | 121 | 0.98 | 122 | 0.98 | 124 |
| P22 | 166 | 0.46 | 345 | 0.97 | 354 | 0.99 | 355 | 0.99 | 355 | 0.99 | 356 | 1 | 357 |
| T04 | 6 | 0.67 | 7 | 0.78 | 9 | 1 | 9 | 1 | 9 | 1 | 9 | 1 | 9 |
| V73 | 194 | 0.99 | 122 | 0.62 | 182 | 0.93 | 194 | 0.99 | 195 | 0.99 | 196 | 1 | 196 |

Legend:

Count: Number of cells that properly classify

Prop: Proportion of cells that properly classify

Total: Total cell count for the donor population
