## Supplemental Table 6 for "Comparison of three quantitative approaches for estimating time-since-deposition from autofluorescence and morphological profiles of cell populations from forensic biological samples"

**Supplemental Table S6: Classifier performance for each hold-out donor/timepoint cell population, using GBM models.**

|  | **Under 7 days** | | **Under 30 days** | | **Under 60 days** | | **Under 90 days** | | **Under 120 days** | | **Under 180 days** | |  |
| --- | --- | --- | --- | --- | --- | --- | --- | --- | --- | --- | --- | --- | --- |
| **Donor** | *Count* | *Prop* | *Count* | *Prop* | *Count* | *Prop* | *Count* | *Prop* | *Count* | *Prop* | *Count* | *Prop* | **Total** |
| A24.125 | 75 | 1 | 65 | 0.87 | 28 | 0.37 | 7 | 0.09 | 5 | 0.07 | 71 | 0.95 | 75 |
| A24.158 | 46 | 1 | 23 | 0.5 | 8 | 0.17 | 2 | 0.04 | 2 | 0.04 | 45 | 0.98 | 46 |
| A24.22 | 103 | 1 | 62 | 0.6 | 98 | 0.95 | 103 | 1 | 103 | 1 | 103 | 1 | 103 |
| A24.226 | 23 | 0.92 | 12 | 0.48 | 1 | 0.04 | 0 | 0 | 0 | 0 | 0 | 0 | 25 |
| A24.414 | 31 | 1 | 28 | 0.9 | 23 | 0.74 | 18 | 0.58 | 14 | 0.45 | 10 | 0.32 | 31 |
| A24.75 | 6 | 1 | 4 | 0.67 | 2 | 0.33 | 5 | 0.83 | 5 | 0.83 | 6 | 1 | 6 |
| A24.83 | 11 | 0.85 | 8 | 0.62 | 3 | 0.23 | 10 | 0.77 | 11 | 0.85 | 13 | 1 | 13 |
| B32.22 | 70 | 0.95 | 42 | 0.57 | 70 | 0.95 | 74 | 1 | 74 | 1 | 74 | 1 | 74 |
| B32.37 | 571 | 1 | 300 | 0.52 | 511 | 0.89 | 550 | 0.96 | 562 | 0.98 | 568 | 0.99 | 572 |
| B32.72 | 8 | 0.89 | 8 | 0.89 | 6 | 0.67 | 5 | 0.56 | 6 | 0.67 | 8 | 0.89 | 9 |
| B32.8 | 20 | 0.95 | 11 | 0.52 | 20 | 0.95 | 21 | 1 | 21 | 1 | 21 | 1 | 21 |
| C58.1 | 146 | 0.83 | 175 | 1 | 175 | 1 | 175 | 1 | 175 | 1 | 175 | 1 | 175 |
| C58.4 | 186 | 0.7 | 263 | 1 | 264 | 1 | 264 | 1 | 264 | 1 | 264 | 1 | 264 |
| D68.3 | 1 | 0.5 | 1 | 0.5 | 2 | 1 | 2 | 1 | 2 | 1 | 2 | 1 | 2 |
| E10.4 | 18 | 0.42 | 41 | 0.95 | 43 | 1 | 43 | 1 | 43 | 1 | 43 | 1 | 43 |
| H21.10 | 43 | 0.98 | 26 | 0.59 | 39 | 0.89 | 42 | 0.95 | 43 | 0.98 | 43 | 0.98 | 44 |
| H21.31 | 194 | 0.9 | 51 | 0.24 | 206 | 0.95 | 213 | 0.99 | 215 | 1 | 215 | 1 | 216 |
| H21.8 | 14 | 1 | 8 | 0.57 | 13 | 0.93 | 14 | 1 | 14 | 1 | 14 | 1 | 14 |
| I66.121 | 169 | 1 | 168 | 0.99 | 144 | 0.85 | 109 | 0.64 | 80 | 0.47 | 130 | 0.77 | 169 |
| I66.158 | 384 | 1 | 366 | 0.95 | 285 | 0.74 | 213 | 0.55 | 163 | 0.42 | 296 | 0.77 | 384 |
| I66.190 | 58 | 0.94 | 27 | 0.44 | 11 | 0.18 | 6 | 0.1 | 3 | 0.05 | 1 | 0.02 | 62 |
| I66.3 | 9 | 0.75 | 12 | 1 | 12 | 1 | 12 | 1 | 12 | 1 | 12 | 1 | 12 |
| I66.333 | 20 | 1 | 20 | 1 | 20 | 1 | 17 | 0.85 | 10 | 0.5 | 4 | 0.2 | 20 |
| I66.35 | 30 | 1 | 27 | 0.9 | 21 | 0.7 | 26 | 0.87 | 28 | 0.93 | 30 | 1 | 30 |
| I66.415 | 114 | 1 | 114 | 1 | 111 | 0.97 | 97 | 0.85 | 86 | 0.75 | 55 | 0.48 | 114 |
| I66.8 | 59 | 0.98 | 39 | 0.65 | 56 | 0.93 | 59 | 0.98 | 60 | 1 | 60 | 1 | 60 |
| I66.83 | 82 | 1 | 77 | 0.94 | 57 | 0.7 | 49 | 0.6 | 68 | 0.83 | 77 | 0.94 | 82 |
| J72.22 | 145 | 0.95 | 100 | 0.65 | 147 | 0.96 | 153 | 1 | 153 | 1 | 153 | 1 | 153 |
| J72.35 | 564 | 1 | 284 | 0.5 | 518 | 0.92 | 554 | 0.98 | 560 | 0.99 | 565 | 1 | 565 |
| J72.8 | 563 | 0.98 | 238 | 0.42 | 524 | 0.92 | 567 | 0.99 | 571 | 1 | 572 | 1 | 572 |
| K47.29 | 53 | 1 | 42 | 0.79 | 52 | 0.98 | 52 | 0.98 | 53 | 1 | 53 | 1 | 53 |
| K47.75 | 41 | 1 | 37 | 0.9 | 24 | 0.59 | 28 | 0.68 | 38 | 0.93 | 41 | 1 | 41 |
| L12.121 | 183 | 1 | 173 | 0.95 | 115 | 0.63 | 72 | 0.39 | 43 | 0.23 | 167 | 0.91 | 183 |
| L12.152 | 572 | 1 | 560 | 0.98 | 443 | 0.77 | 302 | 0.53 | 176 | 0.31 | 514 | 0.9 | 572 |
| L12.22 | 236 | 0.95 | 194 | 0.78 | 244 | 0.98 | 248 | 1 | 248 | 1 | 248 | 1 | 249 |
| L12.353 | 4 | 1 | 4 | 1 | 4 | 1 | 2 | 0.5 | 2 | 0.5 | 0 | 0 | 4 |
| L12.47 | 38 | 0.97 | 18 | 0.46 | 33 | 0.85 | 38 | 0.97 | 39 | 1 | 39 | 1 | 39 |
| L12.59 | 27 | 0.96 | 11 | 0.39 | 26 | 0.93 | 28 | 1 | 28 | 1 | 28 | 1 | 28 |
| L12.82 | 557 | 0.97 | 180 | 0.31 | 26 | 0.05 | 563 | 0.98 | 566 | 0.99 | 570 | 1 | 572 |
| L49.1 | 12 | 0.39 | 28 | 0.9 | 30 | 0.97 | 30 | 0.97 | 30 | 0.97 | 30 | 0.97 | 31 |
| N90.1 | 23 | 0.19 | 108 | 0.87 | 121 | 0.98 | 121 | 0.98 | 122 | 0.98 | 122 | 0.98 | 124 |
| P22.1 | 126 | 0.86 | 147 | 1 | 147 | 1 | 147 | 1 | 147 | 1 | 147 | 1 | 147 |
| P22.3 | 51 | 0.77 | 65 | 0.98 | 65 | 0.98 | 65 | 0.98 | 65 | 0.98 | 65 | 0.98 | 66 |
| P22.4 | 94 | 0.65 | 142 | 0.99 | 144 | 1 | 144 | 1 | 144 | 1 | 144 | 1 | 144 |
| T04.3 | 6 | 0.67 | 8 | 0.89 | 9 | 1 | 9 | 1 | 9 | 1 | 9 | 1 | 9 |
| V73.47 | 146 | 0.99 | 102 | 0.69 | 119 | 0.8 | 141 | 0.95 | 146 | 0.99 | 147 | 0.99 | 148 |
| V73.8 | 46 | 0.96 | 32 | 0.67 | 44 | 0.92 | 47 | 0.98 | 48 | 1 | 48 | 1 | 48 |

Legend:

Count: Number of cells that properly classify

Prop: Proportion of cells that properly classify

Total: Total cell count for the donor population
