## Supplemental Table 7 for "Comparison of three quantitative approaches for estimating time-since-deposition from autofluorescence and morphological profiles of cell populations from forensic biological samples"

**Supplemental Table S7: Classifier performance for each hold-out donor/timepoint cell population, using GLMM models.**

|  | **Under 7 days** | | **Under 30 days** | | **Under 60 days** | | **Under 90 days** | | **Under 120 days** | | **Under 180 days** | |  |
| --- | --- | --- | --- | --- | --- | --- | --- | --- | --- | --- | --- | --- | --- |
| **Donor** | *Count* | *Prop* | *Count* | *Prop* | *Count* | *Prop* | *Count* | *Prop* | *Count* | *Prop* | *Count* | *Prop* | **Total** |
| A24.125 | 75 | 1 | 62 | 0.83 | 18 | 0.24 | 7 | 0.09 | 4 | 0.05 | 74 | 0.99 | 75 |
| A24.158 | 46 | 1 | 25 | 0.54 | 4 | 0.09 | 2 | 0.04 | 1 | 0.02 | 46 | 1 | 46 |
| A24.22 | 103 | 1 | 56 | 0.54 | 97 | 0.94 | 103 | 1 | 103 | 1 | 103 | 1 | 103 |
| A24.226 | 23 | 0.92 | 9 | 0.36 | 0 | 0 | 0 | 0 | 0 | 0 | 0 | 0 | 25 |
| A24.414 | 30 | 0.97 | 26 | 0.84 | 19 | 0.61 | 11 | 0.35 | 9 | 0.29 | 5 | 0.16 | 31 |
| A24.75 | 6 | 1 | 4 | 0.67 | 2 | 0.33 | 5 | 0.83 | 5 | 0.83 | 5 | 0.83 | 6 |
| A24.83 | 12 | 0.92 | 9 | 0.69 | 2 | 0.15 | 11 | 0.85 | 12 | 0.92 | 13 | 1 | 13 |
| B32.22 | 72 | 0.97 | 41 | 0.55 | 73 | 0.99 | 73 | 0.99 | 74 | 1 | 74 | 1 | 74 |
| B32.37 | 571 | 1 | 297 | 0.52 | 527 | 0.92 | 561 | 0.98 | 567 | 0.99 | 571 | 1 | 572 |
| B32.72 | 8 | 0.89 | 8 | 0.89 | 5 | 0.56 | 7 | 0.78 | 9 | 1 | 9 | 1 | 9 |
| B32.8 | 20 | 0.95 | 15 | 0.71 | 20 | 0.95 | 21 | 1 | 21 | 1 | 21 | 1 | 21 |
| C58.1 | 88 | 0.5 | 175 | 1 | 175 | 1 | 175 | 1 | 175 | 1 | 175 | 1 | 175 |
| C58.4 | 108 | 0.41 | 258 | 0.98 | 263 | 1 | 263 | 1 | 263 | 1 | 264 | 1 | 264 |
| D68.3 | 1 | 0.5 | 1 | 0.5 | 2 | 1 | 2 | 1 | 2 | 1 | 2 | 1 | 2 |
| E10.4 | 10 | 0.23 | 36 | 0.84 | 42 | 0.98 | 43 | 1 | 43 | 1 | 43 | 1 | 43 |
| H21.10 | 43 | 0.98 | 27 | 0.61 | 40 | 0.91 | 43 | 0.98 | 43 | 0.98 | 43 | 0.98 | 44 |
| H21.31 | 208 | 0.96 | 53 | 0.25 | 209 | 0.97 | 213 | 0.99 | 215 | 1 | 216 | 1 | 216 |
| H21.8 | 14 | 1 | 10 | 0.71 | 14 | 1 | 14 | 1 | 14 | 1 | 14 | 1 | 14 |
| I66.121 | 169 | 1 | 164 | 0.97 | 142 | 0.84 | 103 | 0.61 | 74 | 0.44 | 129 | 0.76 | 169 |
| I66.158 | 383 | 1 | 363 | 0.95 | 262 | 0.68 | 164 | 0.43 | 103 | 0.27 | 336 | 0.88 | 384 |
| I66.190 | 60 | 0.97 | 32 | 0.52 | 17 | 0.27 | 5 | 0.08 | 4 | 0.06 | 1 | 0.02 | 62 |
| I66.3 | 9 | 0.75 | 12 | 1 | 12 | 1 | 12 | 1 | 12 | 1 | 12 | 1 | 12 |
| I66.333 | 20 | 1 | 20 | 1 | 20 | 1 | 15 | 0.75 | 8 | 0.4 | 5 | 0.25 | 20 |
| I66.35 | 30 | 1 | 23 | 0.77 | 23 | 0.77 | 29 | 0.97 | 30 | 1 | 30 | 1 | 30 |
| I66.415 | 114 | 1 | 114 | 1 | 111 | 0.97 | 99 | 0.87 | 86 | 0.75 | 61 | 0.54 | 114 |
| I66.8 | 59 | 0.98 | 42 | 0.7 | 56 | 0.93 | 59 | 0.98 | 60 | 1 | 60 | 1 | 60 |
| I66.83 | 82 | 1 | 74 | 0.9 | 37 | 0.45 | 70 | 0.85 | 79 | 0.96 | 79 | 0.96 | 82 |
| J72.22 | 150 | 0.98 | 93 | 0.61 | 148 | 0.97 | 152 | 0.99 | 152 | 0.99 | 152 | 0.99 | 153 |
| J72.35 | 564 | 1 | 239 | 0.42 | 542 | 0.96 | 559 | 0.99 | 561 | 0.99 | 565 | 1 | 565 |
| J72.8 | 565 | 0.99 | 261 | 0.46 | 543 | 0.95 | 569 | 0.99 | 572 | 1 | 572 | 1 | 572 |
| K47.29 | 53 | 1 | 41 | 0.77 | 52 | 0.98 | 52 | 0.98 | 52 | 0.98 | 53 | 1 | 53 |
| K47.75 | 41 | 1 | 33 | 0.8 | 18 | 0.44 | 35 | 0.85 | 40 | 0.98 | 40 | 0.98 | 41 |
| L12.121 | 183 | 1 | 163 | 0.89 | 102 | 0.56 | 58 | 0.32 | 37 | 0.2 | 164 | 0.9 | 183 |
| L12.152 | 572 | 1 | 548 | 0.96 | 330 | 0.58 | 150 | 0.26 | 72 | 0.13 | 553 | 0.97 | 572 |
| L12.22 | 244 | 0.98 | 203 | 0.82 | 245 | 0.98 | 248 | 1 | 248 | 1 | 249 | 1 | 249 |
| L12.353 | 4 | 1 | 4 | 1 | 3 | 0.75 | 2 | 0.5 | 2 | 0.5 | 0 | 0 | 4 |
| L12.47 | 38 | 0.97 | 16 | 0.41 | 36 | 0.92 | 39 | 1 | 39 | 1 | 39 | 1 | 39 |
| L12.59 | 27 | 0.96 | 6 | 0.21 | 28 | 1 | 28 | 1 | 28 | 1 | 28 | 1 | 28 |
| L12.82 | 557 | 0.97 | 112 | 0.2 | 13 | 0.02 | 568 | 0.99 | 569 | 0.99 | 571 | 1 | 572 |
| L49.1 | 9 | 0.29 | 27 | 0.87 | 31 | 1 | 31 | 1 | 31 | 1 | 31 | 1 | 31 |
| N90.1 | 16 | 0.13 | 109 | 0.88 | 116 | 0.94 | 121 | 0.98 | 121 | 0.98 | 122 | 0.98 | 124 |
| P22.1 | 75 | 0.51 | 145 | 0.99 | 147 | 1 | 147 | 1 | 147 | 1 | 147 | 1 | 147 |
| P22.3 | 38 | 0.58 | 64 | 0.97 | 65 | 0.98 | 65 | 0.98 | 65 | 0.98 | 65 | 0.98 | 66 |
| P22.4 | 53 | 0.37 | 136 | 0.94 | 142 | 0.99 | 143 | 0.99 | 143 | 0.99 | 144 | 1 | 144 |
| T04.3 | 6 | 0.67 | 7 | 0.78 | 9 | 1 | 9 | 1 | 9 | 1 | 9 | 1 | 9 |
| V73.47 | 147 | 0.99 | 88 | 0.59 | 134 | 0.91 | 146 | 0.99 | 147 | 0.99 | 148 | 1 | 148 |
| V73.8 | 47 | 0.98 | 34 | 0.71 | 48 | 1 | 48 | 1 | 48 | 1 | 48 | 1 | 48 |

Legend:

Count: Number of cells that properly classify

Prop: Proportion of cells that properly classify

Total: Total cell count for the donor population
